# How the AFF1/4 scaffold recruits the elongation factor ELL2 to promote HIV-1 proviral transcription

**DOI:** 10.1101/068262

**Authors:** Shiqian Qi, Zichong Li, Ursula Schulze-Gahmen, Goran Stjepanovic, Qiang Zhou, James H. Hurley

**Affiliations:** State Key Laboratory of Biotherapy, West China Hospital, Sichuan University, and Collaborative Innovation Center for Biotherapy, Chengdu, China; Department of Molecular and Cell Biology and California Institute of Quantitative Biosciences, University of California, Berkeley, Berkeley, United States; Molecular Biophysics and Integrated Bioimaging Division, Lawrence Berkeley National Laboratory, Berkeley, United States

## Abstract

The intrinsically disordered scaffold proteins AFF1/4 and the transcription elongation factors ELL1/2 are core components of the superelongation complex required for HIV-1 proviral transcription. We determined the 2.0-Å resolution crystal structure of the human ELL2 C-terminal domain bound to its 50-residue binding site on AFF4, the ELLBow. The ELL2 domain has the same arch-shaped fold as the tight junction protein occludin. The ELLBow consists of an N-terminal helix followed by an extended hairpin that we refer to as the elbow joint, and occupies most of the concave surface of ELL2. This surface is important for the ability of ELL2 to promote HIV-1 Tat-mediated proviral transcription. The AFF4-ELL2 interface is imperfectly packed, leaving a cavity suggestive of a potential binding site for transcription-promoting small molecules.

## Introduction

Curing Acquired Immunodeficiency Syndrome (AIDS) is a major global health goal. AIDS is caused by the human immunodeficiency virus (HIV), which has proved exceptionally difficult to eradicate (Archin et al., 2014). The principal obstacle to HIV eradication is the persistence in patients of a reservoir of cells harboring latent provirus integrated within the genome (Ruelas and Greene, 2013). Clinical interest in the reactivation of latent HIV (Archin and Margolis, 2014) has brought renewed attention to the mechanism of transcriptional regulation of the HIV provirus. Latency is regulated at the levels of epigenetic silencing and transcription initiation and elongation (Mbonye and Karn, 2014). Transcription elongation, which is promoted by the HIV Tat protein and TAR RNA sequence, is the best-understood of these mechanisms. The functions of HIV Tat and TAR in promoting elongation are completely dependent on their ability to hijack the host Super Elongation Complex (SEC) (He et al., 2010; Lu et al., 2014; Sobhian et al., 2010).

The SEC consists of the Cyclin-dependent kinase CDK9 and Cyclin T (CycT1 or T2), together known as P-TEFb (Price, 2000); one of either of the intrinsically disordered (ID) scaffold proteins AFF1 or AFF4 (He et al., 2010; Sobhian et al., 2010); one of either ENL or AF9; and one of either of the RNA polymerase elongation factors ELL1 or ELL2 (Biswas et al., 2011; He et al., 2010; Luo et al., 2012). The reason that Tat is such a powerful activator of HIV-1 transcription lies in its ability to pack two distinct transcriptional elongation factors P-TEFb and ELL1/2 into a single SEC complex, where the two factors can synergistically stimulate a single RNA Pol II elongation complex (He et al., 2010; Lu et al., 2014). AFF1/4 is more than 1100 residues long and is the principal scaffold that holds the SEC together (Lin et al., 2010). AFF1/4 consists almost entirely of predicted intrinsically disordered regions (IDRs). AFF1 and AFF4 function in transcription elongation by virtue of various peptide motifs interspersed throughout their sequences, much like many other ID signaling and regulatory proteins that have come under intensive study (Csizmok et al., 2016; Tompa et al., 2014). The AFF1- and ELL2-containing version of the SEC is the most important in the promotion of proviral elongation, despite its low abundance (Li et al., 2016).

The structure of P-TEFb lacking the C-terminal IDR of CycT1 has been determined in complex with HIV-1 Tat (Tahirov et al., 2010) and the N-terminal 60 residues of AFF4 (Gu et al., 2014; Schulze-Gahmen et al., 2013). This structure shows that AFF4 residues 32-67 bind as an extended strand followed by two α-helices to the CycT1 surface. NMR studies showed that AFF4 residues 761-774 fold into a β-strand that combines with two strands of the AF9 AFF4-binding domain to generate a three-stranded β-sheet (Leach et al., 2013). The structures of the P-TEFb and AF9 complexes with AFF4 revealed two of the three known interfaces used by AFF4 in assembly of the SEC. In this study, we set out to visualize the last of the three known interfaces critical for AFF4 function, its binding site for ELL1/2.

Progress in characterizing the AFF4 interface with ELL2 has been slower than for the P-TEFb and ENL/AF9 interfaces, in part because the AFF4 binding domain for ELL2 is poorly soluble and prone to aggregation. The first step in this study was to obtain a fusion construct such that a stable obligate complex between ELL2 and AFF4 was formed. This fusion-based tethered complex was stable and soluble enough to be crystallized. The crystal structure confirmed that the AFF4 binding domain of ELL2 has an occludin fold, as predicted from sequence homology (Li et al., 2005). It showed that the IDR consisting of AFF4 residues 301-351 (hereafter referred to as AFF4^ELLBow^ for ELL1/2 Binding) folds up into a helix and elbow joint arrangement that makes extensive contacts with the occludin domain of ELL2 (hereafter ELL2^Occ^). These results complete the structural picture of how AFF1/4 engages its three known partners in the SEC.

## Results

### Mapping the AFF4^ELLBow^ and ELL2^Occ^ interaction

Following the initial mapping of the AFF4 and ELL2 interaction sites to approximately residues 318-337 of the former and 519-640 of the latter (Chou et al., 2013) (Fig. 1A), we sought to isolate a stable form of this monomeric (Fig. 1-S1A) complex for crystallization. It was difficult to obtain diffraction-quality crystals of ELL2^Occ^ constructs with AFF4^ELLBow^ fragments because of the propensity of the ELL2 fragment to aggregate over time. We reasoned that fusion of AFF4^ELLBow^ and ELL2^Occ^ fragments might protect the AFF4 binding epitope on ELL2^Occ^ from aggregation. Constructs were generated for both AFF4^ELLBow^–(Gly-Ser)_4_-ELL2^Occ^ and ELL2^Occ^–(Gly-Ser)_4_-AFF4^ELLBow^. The ELL2^Occ^– (Gly-Ser)_4_-AFF4^ELLBow^ dimerized in solution, while AFF4^ELLBow^–(Gly-Ser)_4_-ELL2^Occ^ was monomeric (Fig. 1-S1B). Given that the unfused fragments were monomeric, we concluded that the dimerization of ELL2^Occ^–(Gly-Ser)_4_- AFF4^ELLBow^ represented a domain-swapping artifact (Fig. 1- S1C) and focused efforts on AFF4^ELLBow^–(Gly-Ser)_4_- ELL2^Occ^.

**Figure 1.**
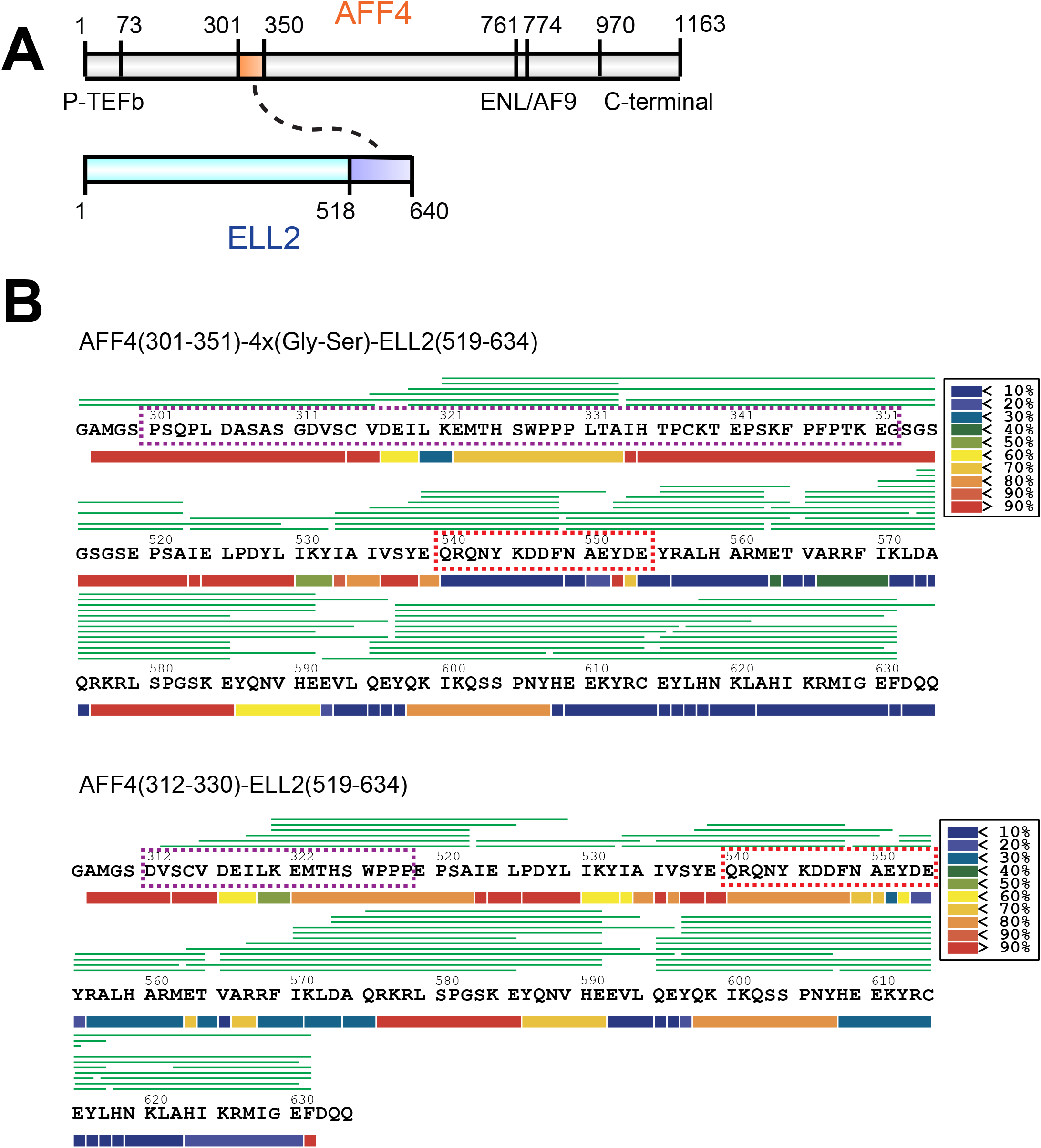
Determinants of AFF4 binding to ELL2. **A.** Schematic of the interactions of the AFF4 IDP scaffold with its partners in the SEC. The highlighted boxes within AFF4 and ELL2 represent the co-crystallized elements described below. Other regions of AFF4 are annotated for binding to P-TEFb, AF9/ENL, and the novel C-terminal ELL1/2 binding site described below. **B**. Deuterium uptake data for the AFF4-ELL2 fusion complexes. HDX-MS data are shown in heat map format, where peptides are represented using rectangular strips above the protein sequence. Absolute deuterium uptake (in %) after 10 s is indicated by a color gradient below the protein sequence. Protein sequence corresponding to the AFF4 is indicated by purple box. Protein sequence corresponding to the N-terminal portion of the ELL2 α1 helix is indicated by red box.

Hydrogen-deuterium exchange coupled to mass spectrometry (HDX-MS) was used to probe the protection of expected helical regions of ELL2^Occ^ in variant constructs consisting of AFF4^301-351^-(Gly-Ser)_4_-ELL2^519-640^ and AFF4^312-330^- ELL2^519-640^. Essentially complete peptide coverage was obtained for both constructs (Fig. 1B). The longer construct manifested some exchange-protected regions (330-332) absent in the shorter construct (see first dashed box above and below in Fig. 1B). The data for the longer construct also showed that the ELL2 sequence 540-554, corresponding to part of the first predicted helix (second dashed box above and below in Fig. 1B), manifested < 10 % exchange over 10 s. In comparison, a higher level of exchange was observed in the shorter construct. We proceeded to focus on crystallizing AFF4^301-351^-(Gly-Ser)_4_-ELL2^519-640^.

### Structure of the AFF4^ELLBow^:ELL2^Occ^ complex

The structure of the AFF4 ^ELLBow^:ELL2^Occ^ complex was determined by SeMet MAD phasing (Fig. 2A, Fig. 2-S1, Table 1). Helix α1 (residues 538-578) bends inward at Tyr552 by 30° such that the C-terminal helix of α1 (553-578) pack against α2 (Fig. 2B). Helices α2 (584-602) and α3 (607-638) of ELL2 are oriented at an angle of ∼100° with respect to each other such that both pack along the length of the long, bent helix α1 (Fig.2B). The structure confirms that ELL2^Occ^ has the same arch-shaped three-helix fold as the C-terminal domain of occludin (Li et al., 2005). The ELL2^Occ^ and occludin C-terminal domain (pdb entry 1XAW) structures can be superimposed with an r.m.s.d. of 4.0 Å for 104 residue pairs (Fig. 2C). The main differences are in the α2-α3 connector and in the mutual orientation of these two helices. The α2-α3 angle is steeper in ELL2^Occ^ than in occludin. A minor difference is that ELL2^Occ^ has an extra single-turn helix, denoted α0, at its N-terminus.

**Figure 2.**
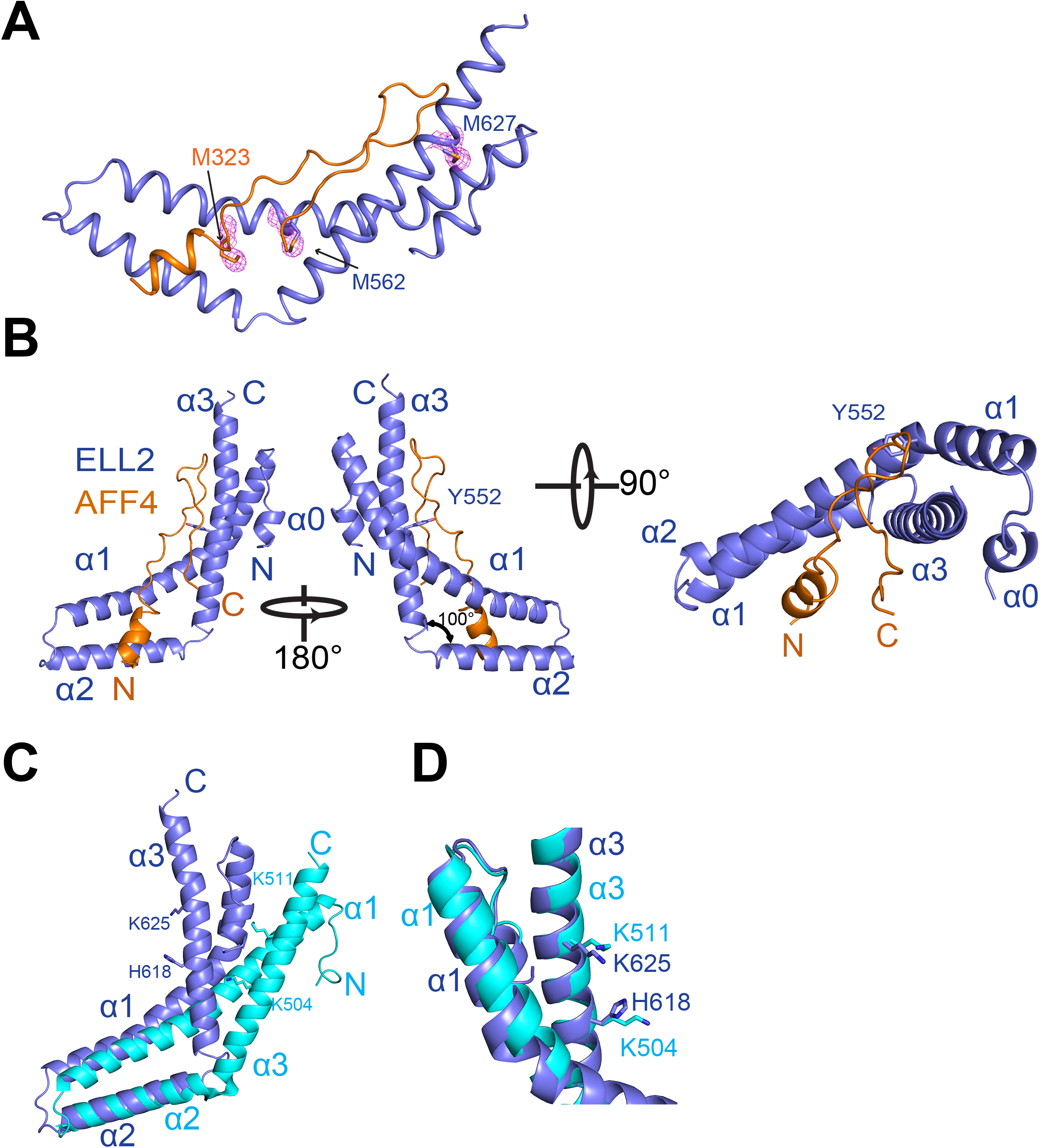
Crystal structure of the AFF4 ELLBow in complex with the Occludin homology domain of ELL2. **A.** Se anomalous difference peaks and overall structure of the complex. The Se substructure map is displayed at a contour level of 2σ(magenta). B. Three views of the overall structure of the complex, with AFF4 in orange and ELL2 in light blue. C. Comparison of ELL2 and occludin C-terminal domain showing that the folds are similar but ELL2 is more sharply bent. D. α3 from both ELL2 and Occludin are aligned, the structurally and functionally conserved residues are shown in stick. ELL2 is shown in light blue while Occludin is shown in cyan.

**Table 1.**
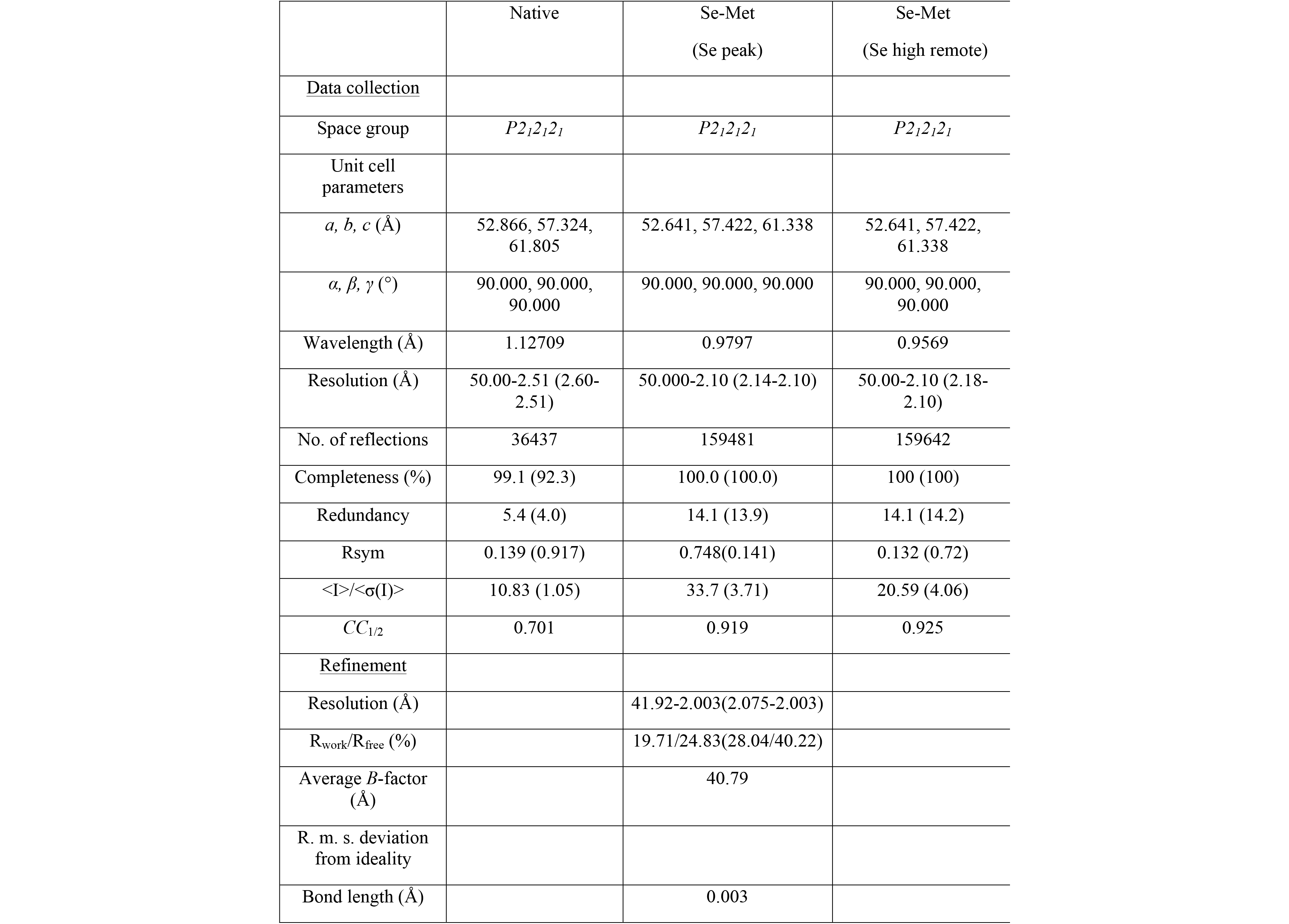
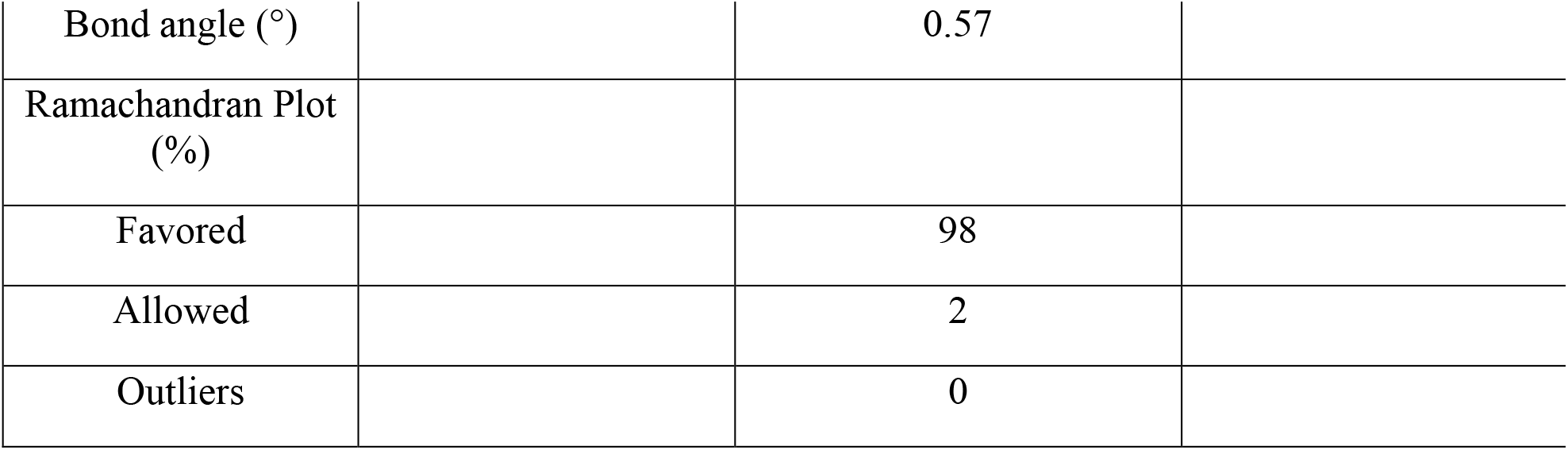
Statistics of Crystallographic Data Reduction and Refinement

AFF4^ELLBow^ is ordered over residues 314-349 and buries 1535 Å^2^ of solvent-accessible surface area. Fully 37% of the entire solvent accessible surface area of AFF4 ^ELLBow^ is buried in the contact. The AFF4 ^ELLBow^ sequence folds into several distinct regions. It begins with helix α1 (315-324), is followed by an extended hydrophobic sequence (325-327), a polyproline segment (328-330), an extended region that doubles back on itself in what we refer to as the ELLBow joint (331-343), and a second extended hydrophobic sequence (344-349) (Fig. 3A). The fusion construct contains 17 residues that are not visualized in electron density, AFF4 351-351, 8 Gly-Ser linker residues, and ELL2 519-524, more than adequate to span the 15 Å between AFF4 349 and ELL2 525 in the structure. Hydrophobic side-chains of AFF4 ^ELLBow^ α1, including Val316, Ile319, Leu320, and Met323, are buried in a hydrophobic groove formed by the C-terminal half of ELL2^Occ-^ α1 and α2 (Fig. 3B). These helices of ELL2^Occ^ contribute hydrophobic residues Val565, Phe569, Ile570, Leu572, Aps573, Val589, His590, Tyr596, Leu594 and Ile599 to the AFF4 α1 binding site (Fig. 3B). ELL2^Occ^ buries 1315 Å^2^ of solvent-accessible surface area, corresponding to 15% of its total surface area.

**Figure 3.**
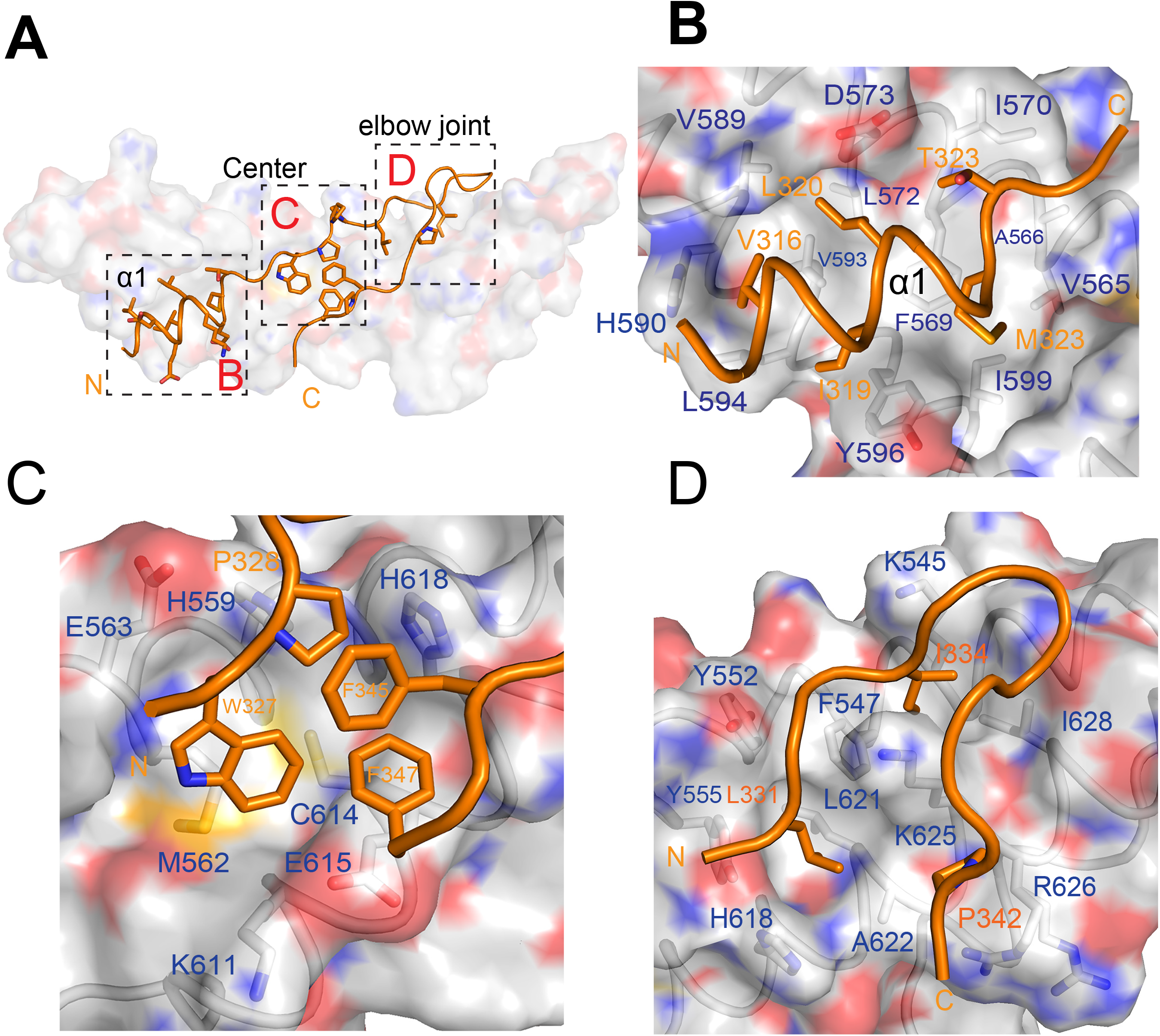
AFF4^ELLBow^:ELL2^Occ^ interaction surfaces. **A.** Overview of the main binding determinants of the AFF4 ELLBow. **B.** The first helix of the ELLBow (orange) binds in a hydrophobic groove on ELL2 (gray). The key residues are shown in a stick model. **C.** The central cluster, in which hydrophobic residues of the ELLBow pack against ELL2 and one another, and are supplemented by polar interactions. Water molecules are shown as red spheres. c **D.** The ELLBow joint.

AFF4 ^ELLBow^ is centered on Trp327, which forms extensive hydrophobic interactions with the side-chains of ELL2 residues His 559, Met562, Cys614 and Glu615. The Trp327 indole nitrogen also forms a water-mediated hydrogen bond with the Tyr607 hydroxyl. This cluster of residues is completed by the side chains of AFF4 Pro328, Phe345 and Phe347 (Fig. 3C). Collectively, this cluster forms an extensive interaction network in which AFF4 ^ELLBow^ folds up not only against ELL2 but also against itself.

In the AFF4 ^ELLBow^ joint, the side chain of Leu331 sticks into a pocket comprising Tyr 552, Tyr555, His618, Leu621 and Ala622 of the N-terminal half of ELL2^Occ-^α1 and α3. The side chain of Ile334 packs against the side chains of Lys545, Phe547, Lys625 and Leu628. At the distal end of ELLBow joint, Pro342 falls into a shallow cavity composed of Ala622, Lys625 and Arg626 (Fig. 3D).

A number of hydrogen bonds are observed in the complex (Fig. 4A). In AFF4 ^ELLBow^ α1, the side-chains of Asp317 and Arg576 of ELL2 form a bidentate salt-bridge with one another (Fig. 4B). Glu322 of AFF4 forms a 2.8 Å salt bridge with one of the two observed rotamers of His608 of ELL2 (Fig. 4B). In the central cluster, the carbonyl group of Pro328 forms a 2.7 Å hydrogen bond with the side chain of His559 of ELL2 (Fig. 4C). Moving into the ELLBow joint, the main chain amide and carbonyl of AFF4 Leu331 form hydrogen bonds with the hydroxyl oxygens of Tyr552 and Tyr555, respectively, of ELL2. A 2.6 Å hydrogen bond is formed between Thr332 of AFF4 and Lys625 of ELL2 (Fig. 4D). The Ile334 carbonyl accepts a hydrogen bond from the side-chain of Lys545. The Cys338 main-chain amide donates a hydrogen bond to the side-chain of Asp632. The main-chain amide of Phe345 forms a 2.9 Å hydrogen bond with the side chain of Gln619 (Fig. 4D).

**Figure 4.**
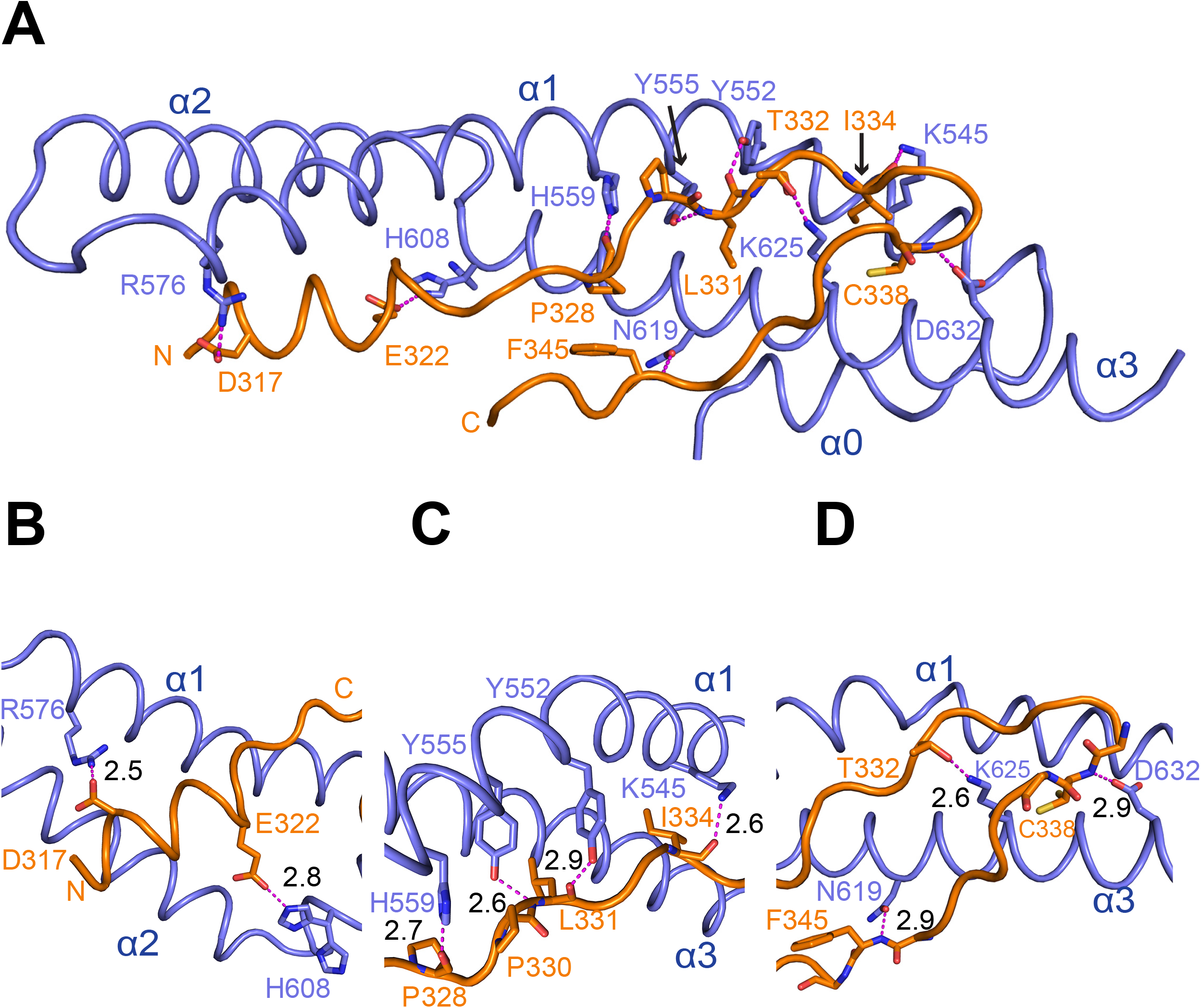
Hydrogen bonding in the AFF4^ELLBow^:ELL2^Occ^ complex. **A.** Overview of the network of hydrogen bonds. The residues involved in the hydrogen bonding are shown in stick. Hydrogen bonds are shown as magenta-colored dashed lines. **(B-D).** Details of the hydrogen bonds. The length of the hydrogen bonds is indicated next to the dashed lines.

The AFF4^ELLBow^:ELL2^Occ^ complex was screened for cavities using POCASA 1.1 (POcket-CAvity Search Application) (Yu et al., 2010) with a probe radius of 3 Å. Of the five largest cavities located, one of these is an internal cavity at the AFF4 ^ELLBow^:ELL2^Occ^ interface (Fig. 5A). The cavity is 36 Å^3^ in volume and is connected to the exterior by a narrow mouth (Fig. 5B). It is lined by the aliphatic part of Glu322, Met323, His325, Trp327, Phe347, and Pro348 of AFF4 and by Met562, Ala566, Tyr607, and the aliphatic part of Lys611 of ELL2 (Fig. 5C). These residues are in or adjoin the central cluster part of the interface.

**Figure 5.**
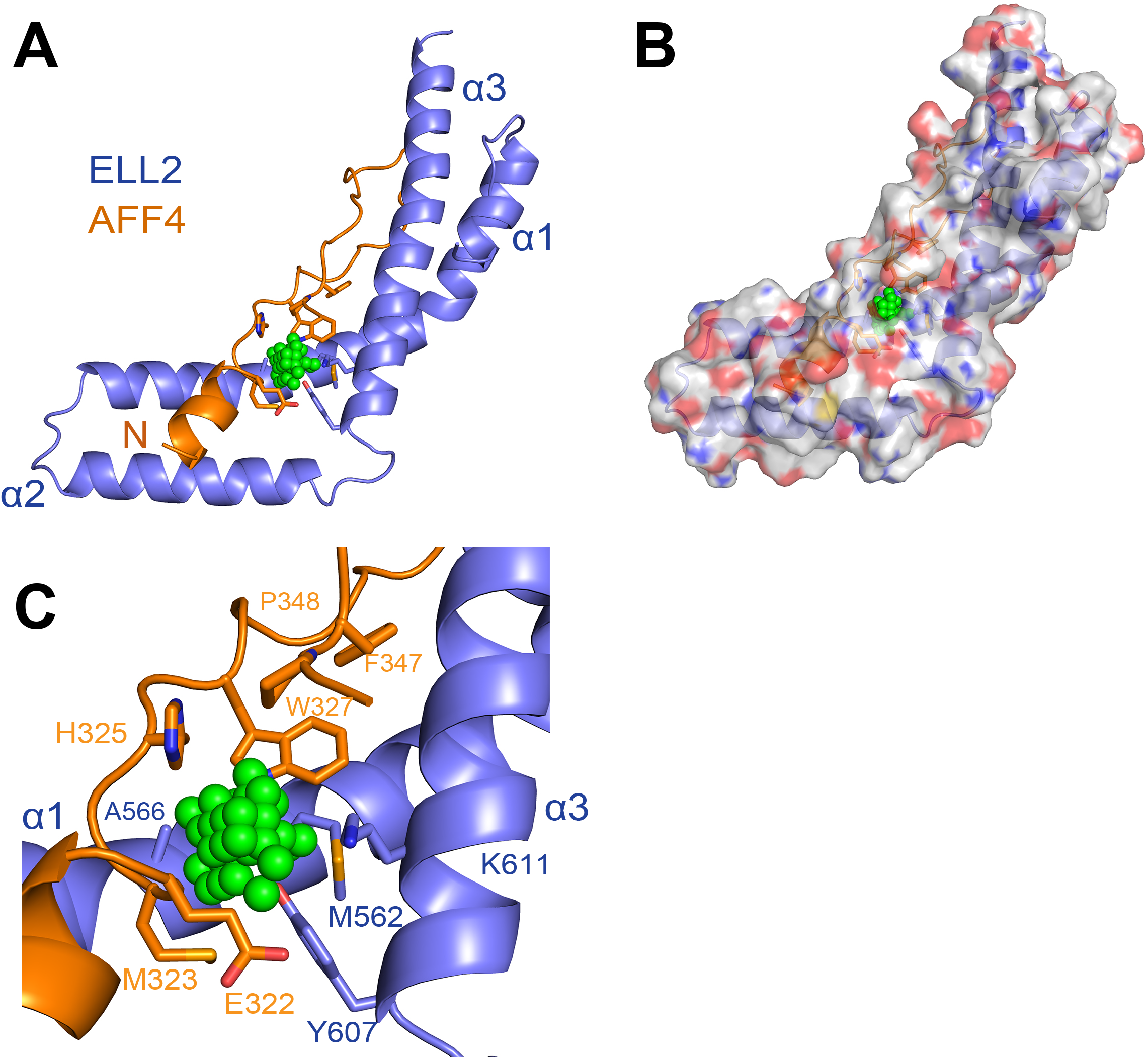
Cavity at the AFF4^ELLBow^:ELL2^Occ^ interface. **A.** Overall view of the cavity that might work as a potential drug binding site. Molecules are shown in cartoon. Side-chains of residues that line the sides of cavity are shown in a stick representation. Colors are light blue, ELL2; orange, AFF4; green, dummy atoms marking the cavity. **B.** Surface model of AFF4^ELLBow^:ELL2^Occ^ complex showing the mouth of the cavity. **C.** Close-up of the cavity region as represented and colored in (A).

### Function of the AFF4 ^ELLBow^:ELL2^Occ^ interface in binding

To validate whether that the observed structural interface corresponded to the mode of binding of AFF4 ^ELLBow^ and ELL2^Occ^ in solution, we carried out a series of mutant peptide binding assays using fluorescence polarization. We considered this particularly critical given the use of the fusion construct to obtain crystals. The assay monitored the displacement of fluorescently labeled wild-type ELLBow peptide by unlabeled mutant peptides 301-351. The unlabeled wild-type peptide in this system has *K_d_* = 86 nM (Table 2; Fig. 6A). The AFF4 hydrophobic residues Val316, Ile319, Leu320, Met323, Trp327, Leu331, Ile334 and Pro342, were mutated to Asp in order to maximally destabilize hydrophobic interactions. Consistent with expectation, mutation of multiple hydrophobic residues to Asp resulted in large decreases in affinity. The double mutant I319D/L320D reduced affinity by >25-fold (Table 2; Fig. 6A). The *K_d_* for the triple mutant I319D/L320D/M323D was immeasurable due to weak binding, but greater than 3 μM, representing a ∼50-fold loss of affinity (Table 2; Fig. 6A). The same was true of two other triple hydrophobic mutants tested, M323D/L331D/I334D and W327D/L331D/I334D (Table 2; Fig. 6B). The single mutant M323D has the largest effect of any single amino acid change, with a reduction in affinity of >25-fold (Table 2; Fig. 6A). Moving closer to the center of the AFF4 ELLBow, L331D and I334D reduce affinity by ∼20- and 8-fold, respectively (Table 2; Fig. 6B). This highlights the role of hydrophobic residues in AFF4 ELLBow helix α1 and immediately C-terminal to it in the central cluster as the critical anchor points and affinity determinants.

**Figure 6.**
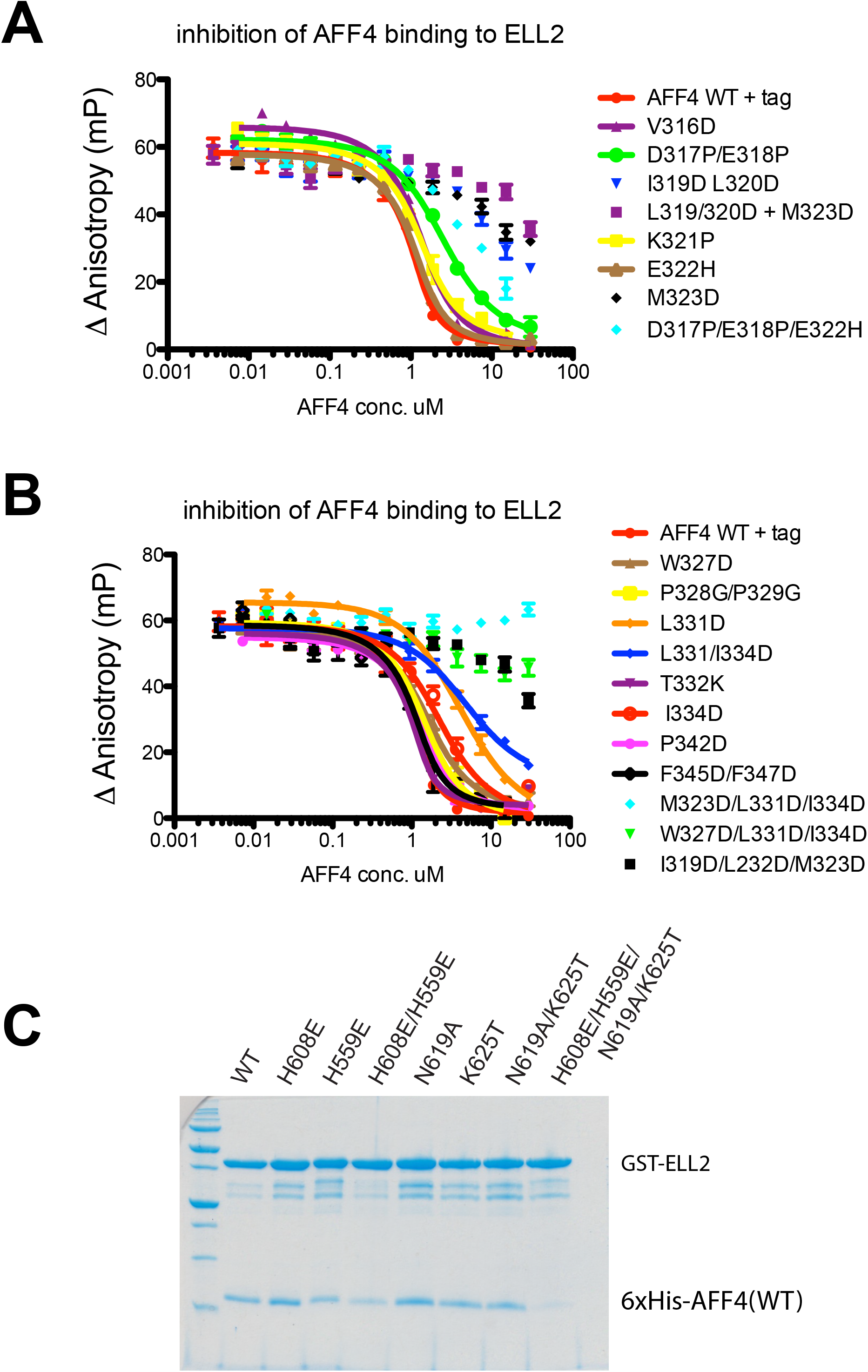
Contributions of AFF4 ELLBow interactions to binding in solution. **A**. Binding of AFF4^ELLBow^ WT and mutants in the N-terminal α-helix to Sumo-ELL2^Occ^. Sumo-ELL2^Occ^ binding to fluorescently labeled AFF1^ELLBow^ is competitively inhibited by increasing amounts of AFF4^ELLBow^, as described in experimental procedures. Error bars reflect the standard error from three experimental replicates. **B.** Binding of AFF4^ELLBow^ WT and mutants in the central cluster and elbow joint, assayed as in (A). **C.** GST-fusions of the indicated ELL2^Occ^ mutants were immobilized and their ability to pull down His_6_-tagged wild-type AFF4^ELLBow^ assessed.

**Table 2.** K_d_ values for AFF4-ELL2 binding as determined by fluorescence anisotropy.

| **AFF4**<br>**construct/mutation** | **$K_d$ (μM)** | **$R^2$** |
| --- | --- | --- |
| WT, 300-350 +His<sub>6</sub>-tag | $0.086 \pm 0.024$ | 0.97 |
| V316D | $0.23 \pm 0.054$ | 0.99 |
| I319D/L320D | NA |  |
| M323D | NA |  |
| D317P/E318P | $0.82 \pm 0.19$ | 0.99 |
| K321P | $0.24 \pm 0.06$ | 0.98 |
| E322H | $0.13 \pm 0.037$ | 0.99 |
| W327D | $0.38 \pm 0.097$ | 0.96 |
| P328G/P329G | $0.28 \pm 0.076$ | 0.97 |
| L331D | $1.6 \pm 0.38$ | 0.98 |
| T332K | $0.069 \pm 0.037$ | 0.97 |
| I334D | $0.69 \pm 0.17$ | 0.95 |
| L331D/I334D | $1.98 \pm 0.58$ | 0.97 |
| P342D | $0.2 \pm 0.056$ | 0.99 |
| F345D/F347D | $0.13 \pm 0.041$ | 0.97 |
| I319D/L232D/M323D | NA |  |
| D317P/E318P/E322H | NA |  |
| M323D/L331D/I334D | NA |  |
| W327D/ L331D/I334D | NA |  |

Hydrophobic residues of the central cluster make smaller contributions than those highlighted above. W327D reduces affinity 4-fold, while F345D/F347D reduces it by less than two-fold. P342D led to a similar 3-fold drop (Table 2; Fig. 6B). These more modest contributions may reflect that these side-chains are partially solvent-accessible in the AFF4 ^ELLBow^:ELL2^Occ^ complex. Moreover, their interactions are made in part with other residues within the AFF4 ^ELLBow^ such that they could potentially make residual hydrophobic interactions even in unbound AFF4. The polyproline helix does not seem to have a major role in affinity, with the double 328-329 Pro-Gly mutant reducing affinity only by a factor of three (Table 2; Fig. 6B).

The interface has a significant polar component, with some hydrophilic residues contributing substantially to binding, and others less so. The AFF4 ^ELLBow^ α1 mutant D317P/E317P was designed to disrupt hydrogen bonding involving Asp317 and to introduce helix breaker mutants in α1. This mutation lowered affinity by 10-fold (Table 2; Fig. 6A). The charge reversal mutation E322H reduced affinity by less than two-fold (Table 2; Fig. 6A).

It proved impossible to purify hydrophobic to Asp mutants in the AFF4 binding site of ELL2^Occ^ because these proteins were insoluble when expressed in *E. coli*. Presumably this is because these hydrophobic residues also contribute to the hydrophobic core of the ELL2^Occ^ fold. It was, however, possible to purify ELL2^Occ^ polar mutants in the binding site. We examined the roles of ELL2 His559, His608, Asn619, and Lys625 by pull-down assay (Fig. 6C). Single mutants H559E, H608E, N619A, and K625T had no apparent effect on binding by pull-down. However, the quadruple mutant H559E/H608E/N619A/K625T completely abrogated binding in this assay. This validates the role of these residues in the interface in solution.

### AFF4^ELLBow^ and ELL2^Occ^ are important for *in vivo* complex assembly

It had previously been shown that the AFF4 sequence 318-337 was sufficient for ELL2 binding (Chou et al., 2013). We tested whether this sequence was also necessary for binding, and found to our surprise that AFF4 Δ318-337 still pulled down ELL2 in nuclear extracts (Fig. 7-S1A). We hypothesized that AFF4 contained a second ELL2 binding site, and determined that a double deletion of residues 318-337 and 970-1163 abrogated the interaction completely (Fig. 7A). In order to determine if single residues within AFF4 ^ELLBow^ contributed to binding and function in cells, point mutants were constructed in the context of AFF4 Δ970-1163. ELL1 contains a C-terminal domain homologous to that of ELL2, hence binding to ELL1 was also tested. L320D was most effective, blocking both ELL1 and ELL2, consistent with its very strong effect on binding *in vitro* (Fig. 6A, Table 2). E322H, P329G, and I334D partially blocked ELL2 binding but completely knocked out ELL1 binding, consistent with their intermediate effects on *in vitro* peptide binding. Both ELL1 and ELL2 bound robustly to the mutants P324D, F345D, and F347D, consistent with their 2-3-fold effects on binding *in vitro* (Fig. 6A, Table 2).

**Figure 7.**
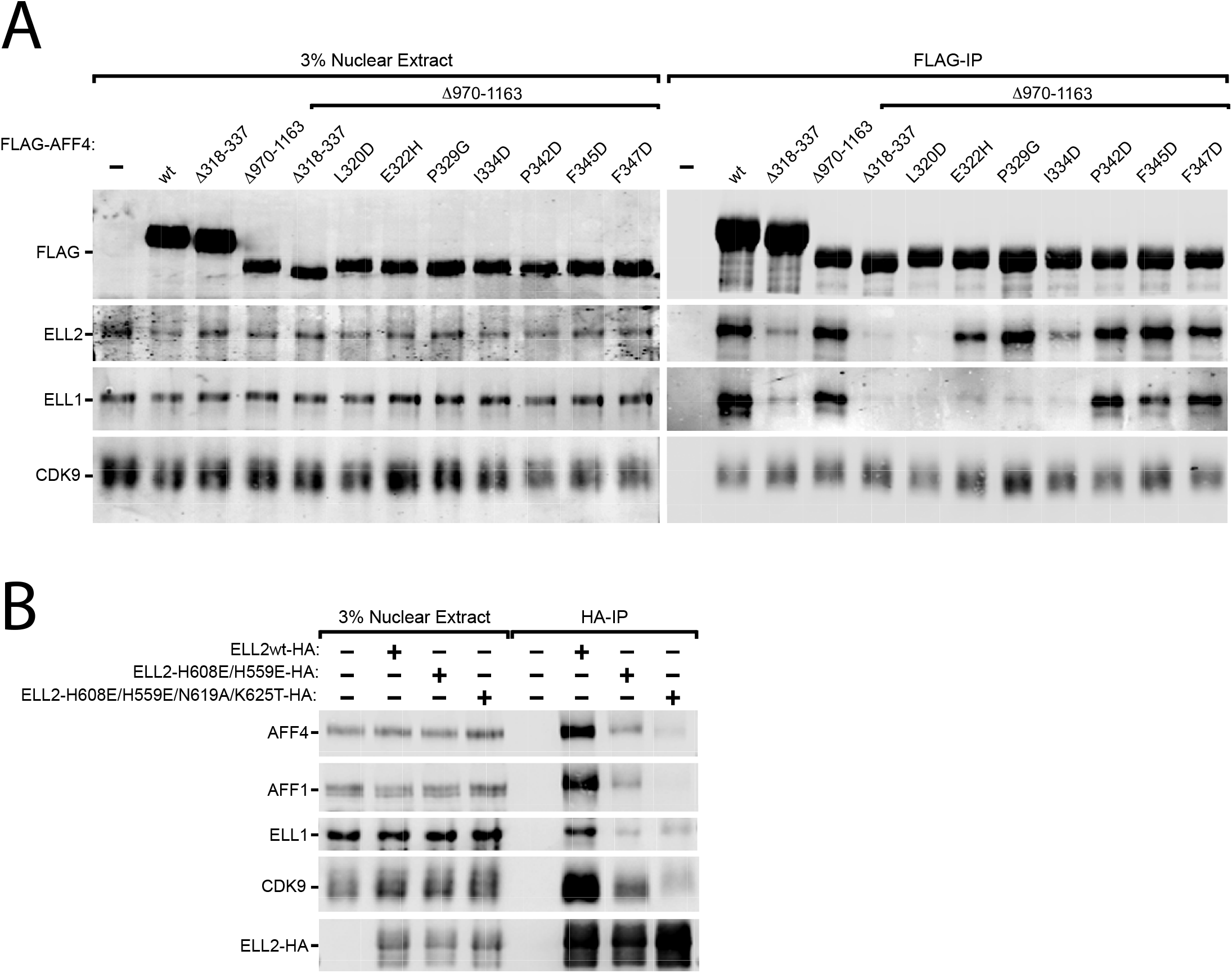
Role of the ELLBow in AFF4 interactions with ELL1/2 in nuclear extracts. Nuclear extracts (NE) were prepared from HEK 293T cells transfected by the indicated plasmids and subjected to immunoprecipitation (IP) with anti-FLAG (**A.**) or anti-HA (**B.**) agarose beads. The NE inputs and IP eluates were examined by immunoblotting for presence of the various proteins labeled on the left.

In order to determine if the AFF4 binding site on ELL2 was functional in cells, polar mutants were inserted into ELL2 alleles and these were transfected into HeLa cells. We avoided testing hydrophobic mutants of ELL2 since we had previously found that these destabilized the ELL2 structure. HA-tagged ELL2^H559E/H608E^ and ELL2^H559E/H608E/N619A/K625T^ were expressed at essentially wild-type levels in HeLa cells (Fig. 7B). Wild-type HA-ELL2 pulled down AFF1, AFF4, and ELL1 from extracts. ELL2^H559E/H608E^ has sharply reduced binding to AFF1, AFF4, and ELL1. ELL2^H559E/H608E/N619A/K625T^ has only trace binding to AFF1 and AFF4 in extracts. These findings support that the structural interface is responsible for the interaction of ELL2 with both AFF1 and AFF4 in cells.

### Role of the ELL2^Occ^ ELLBow-binding interface in HIV proviral transactivation

Overexpression of AFF4 stimulates proviral transcription by ∼5-9-fold and ∼26-fold in HEK 293T and HeLa cells, respectively (Fig. 8A). Unsurprisingly, AFF4 ELLBow mutants that on their own do not abrogate binding have little or no effect on transcription (Fig. 7-S1B). Deletion of the C-terminal ELL1/2 binding domain almost completely blocked transactivation (Fig. 8A). The residual activity of AFF4Δ970-1163 was so low that meaningful results could not be obtained for transactivation phenotypes of these mutants (Fig. 8A). The abundance of the SEC complex appears to be limiting for transactivation such that overexpression of ELL2 in the presence of extra AFF4 promotes transcription by a factor of 14 (Fig. 8B). Polar mutants in the AFF4 binding site of ELL2^Occ^ were tested for their effects on transcription. ELL2^H559E/H608E^ and ELL2^H559E/H608E/N619A/K625T^ had 3-fold and 5-fold less transactivation activity, respectively, than wild-type. These observations strongly support a functional role for the AFF4 ELLBow binding site on ELL2^Occ^ in transactivation.

**Figure 8.**
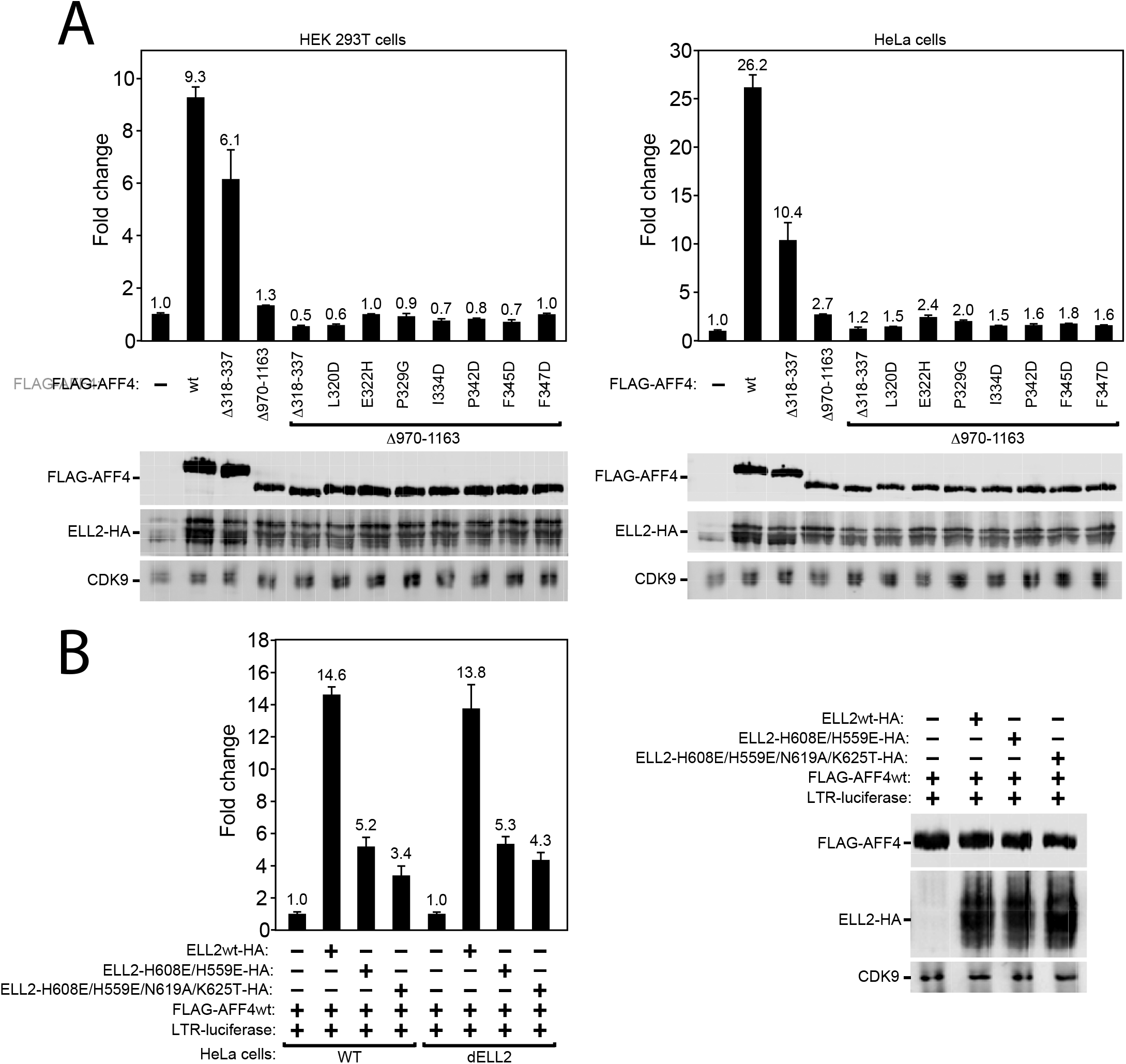
The ELLBow binding site of ELL2 is important for HIV-1 LTR transcription. Luciferase activities were measured and analyzed in extracts of cells transfected in triplicate with the HIV-1 LTR-luciferase construct together with the combinations of plasmids expressing wild-type and mutant ELL2-HA and FLAG-AFF4 as indicated. Each of the ELL2 and AFF4 plasmids was transfected at 1 μg/well. The activities in the control groups were set to 1. The error bars represent mean +/-standard deviation from triplicate wells. An aliquot of each cell extract was examined by immunoblotting for presence of the various proteins labeled on the left.

## Discussion

The crystallization of the AFF4 ^ELLBow^:ELL2^Occ^ complex rounds out our structural-level understanding of how the AFF4 scaffold recruits its three known partners in the SEC, P-TEFb, ENL/AF9, and ELL1/2. The limited solubility of ELL2^Occ^ made this a more challenging target for crystallization, hence the necessity for the fusion approach. When using protein chimeras as a basis for structure solution, it is particularly critical to validate the findings in solution and in functional assays. Binding assays *in vitro*, pull-downs from nuclear extracts, and proviral transactivation assays present a unified, consistent picture that validates the structural results.

The structure confirms the decade-old prediction that the C-terminal domains of ELL1/2 would have the same fold as the occludin ZO-1 binding domain. Occludin is a transmembrane tight junction protein that has no known involvement in transcription. It is not clear why this protein and ELL1/2 should share a domain uniquely present in this small set of otherwise unrelated proteins. In the initial analysis of the occludin structure, it was proposed that another tight junction protein, ZO-1, bound to a basic patch at the concave center of the arch (Li et al., 2005). This patch of occludin includes Lys504 and Lys511, which correspond structurally to the functionally important His618 and Lys625 in the AFF1/4 binding site of ELL2. Subsequently, another report proposed that ZO-1 bound elsewhere, at one tip of the occludin domain arch. Despite these uncertainties, the structural similarities are extensive enough to suggest a common evolutionary origin and related protein-binding functions for the three-helical domains of occludin and ELL1/2.

The bromodomain and extraterminal (BET) protein inhibitor JQ1 (Filippakopoulos et al., 2010) and related compounds promote reactivation of HIV-1 from latency via P-TEFb (Banerjee et al., 2012; Bartholomeeusen et al., 2012; Boehm et al., 2013; Li et al., 2013; Zhu et al., 2012). Other small molecule activators of HIV-1 transcription are being sought in the context of HIV eradication strategies. We observed a cavity at the AFF4-ELL2 interface that appears likely to be present also in the AFF1-ELL2 complex relevant to proviral activation (Li et al., 2016), on the basis of the complete identity of the AFF1 and AFF4 residues involved. If so, this could provide an avenue for the design of new SEC activators with JQ1-like effects on latency, but acting by an orthogonal molecular mechanism.

## Experimental procedures

### Cloning and protein purification

DNAs for ELL2 fragments and AFF4-ELL2 fusions were subcloned into pGST-parallel2, and DNAs for AFF4 peptide fragments were subcloned into pRSFduet-1 and pHis-parallel2. Plasmids expressing FLAG-tagged wild-type AFF4 and HA-tagged wild-type ELL2 were generated previously (He et al., 2010). The plasmids expressing mutant versions of AFF4 and ELL2 were generated by PCR mutagenesis. The mutant constructs were verified by DNA sequencing. All proteins were expressed in *E. coli* BL21-gold (DE3) cells (Agilent Technologies). After induction with 0.2 mM IPTG overnight at 16 °C, the cells were pelleted by centrifugation at 4000 x *g* for 10 minutes. Cell pellets were lysed in 25 mM Tris-HCl pH 8.0, 150 mM NaCl, 0.5 mM TCEP-HCl, and 1 mM PMSF by ultrasonication. The lysate were centrifuged at 25,000 x *g* for 1 hour at 4°C. The supernatants for ELL2 and its fusions were loaded onto GS4B resin at 4°C, target proteins were eluted, and the eluate applied to a Hi Trap Q HP column. Peak fractions were collected and digested with Tobacco Etch Virus (TEV) protease at 4°C overnight. TEV and GST were removed by loading the solution onto Ni-NTA and GS4B columns, respectively. Target proteins were further purified on a Superdex 200 16/60 column equilibrated with 25 mM Tris-HCl pH 8.0, 150 mM NaCl, and 0.5 mM TCEP-HCl. The peak fractions were collected and flash-frozen in liquid N2 for storage. The supernatant of AFF4 was loaded onto Ni-NTA resin at 4°C, eluted with an imidazole gradient, and applied to a Superdex 75 16/60 column equilibrated with 25 mM Tris-HCl pH 8.0, 150 mM NaCl, 0.5 mM TCEP-HCl. Selenomethionyl (SeMet) protein was expressed in *E.coli* BL21-gold (DE3) cells grown in M9 minimal medium supplemented with 5% LB medium. 0.2 mM IPTG and 100 mg selenomethionine were added when the OD_600_ reached 1.0. Cells were pelleted by centrifugation at 4000 x *g* for 10 minutes after overnight induction at 16 °C. Se-Met AFF4^301-351^-(Gly-Ser)_4_-ELL2^519-640^ was prepared as above and SeMet incorporation verified by mass spectrometry.

### HDX-MS Experiments

Amide HDX was initiated by a 20-fold dilution of 40 μM AFF4-ELL2 fusion into a D_2_O buffer containing 25 mM Tris-HCl (pH 8.0), 150 mM NaCl, and 5 mM DTT at 30°C. After 10 s, exchange was quenched at 0°C with the addition of ice-cold quench buffer (400 mMKH2PO_4_/H_3_PO_4_ [pH 2.2]). Quenched samples were injected onto a high-performance liquid chromatography (HPLC) (Agilent 1100; Agilent Technologies) with in-line peptic digestion and desalting. Desalted peptides were eluted and directly analyzed by an Orbitrap Discovery mass spectrometer (Thermo Scientific). Peptides were identified using tandem MS/MS and Proteome Discoverer 2.1 (Thermo Scientific). Initial mass analysis of the peptide centroids was performed using HDExaminer2.0 (Sierra Analytics) followed by manual verification of every peptide. The deuteron content was adjusted for deuteron gain/loss during digestion and HPLC. Both non-deuterated and fully deuterated fusions were analyzed. Fully deuterated samples were prepared by three cycles of drying and resolubilization in D_2_O and 6 M guanidinium hydrochloride.

### Crystallization of the AFF4^ELLBow^-ELL2^Occ^ fusion

The purified fusion construct AFF4(301-351)-(Gly-Ser)4-ELL2(519-640) was concentrated to 10 mg/ml with a 10 kD centrifugal filter (Millipore). Crystals were grown by hanging-drop vapor-diffusion at 19°C. The protein solution was mixed with well buffer composed of 0.2 M NaCl, 10 mM MgCl_2_, 0.3M Na_3_ Citrate, 0.2M Na thiocyanate, 0.1M Hepes pH 7.4. Crystals appeared in 24 hr and grew to full size in 5 days. Crystals were flash-frozen with liquid N_2_ in well buffer. Se-Met crystals were grown in the same condition as native crystals.

### Data collection and structure determination

Native data were collected on BL7-1 at Stanford Synchrotron Radiation Lightsource. Native crystals diffracted to 2.5 Å and data were collected at a wavelength of 1.1271 Å.Se-MAD data were collected on BL8.3.1 at the Advanced Light Source, LBNL, Berkeley. The peak data set and the high energy remote data set were collected at wavelengths of 0.9797Å and 0.9569Å respectively. All data sets were processed with HKL2000 (HKL Research). Data collection and processing statistics are given in Table 1. Phases were determined by two-wavelength multiwavelength anomalous dispersion (MAD). Model building and refinement were finished with ARP/wARP (Langer et al., 2008), COOT (Emsley et al., 2010), REFMAC5 (Murshudov et al., 1997) and PHENIX (Adams et al.,2010).

### Pulldown Assays

Mutants of ELL2 (519-640) and AFF4 (300-351) were purified as described above. The concentration of proteins and peptides was determined by UV absorption at 260-280-nm.9 μM GST-ELL2 and 20 μM His_6_-AFF4 were incubated with GS4B resin at 4°C for 2 hours in 80 μL of 25 mM Tris-HCl pH 8.0, 150 mM NaCl, 0.5mM TCEP-HCl. The resin was washed 3 times with the incubation buffer. Then, the resin was boiled in 30 μL 1x SDS loading buffer at 95°C for 5 min before being applied to SDS-PAGE for analysis.

### Fluorescence Polarization

Protein binding was measured using the fluorescence anisotropy of a 33-residue segment of AFF1 (residues 358-390) encompassing the protein-protein contacts in the crystal structure. AFF1 358-390 are almost identical to AFF4 318-350 with only 3 amino acid changes between the two homologs. The AFF1 peptide was synthesized at the University of Utah DNA/Peptide Facility using the following sequence: C-FAM-GABA-EILKEMTHSWPPPLTAIHTPSTAEPSKFPFPTK-amide where FAM indicates 5-carboxyfluoroscein and GABA indicates a γ-amino-butyric acid spacer. Increasing amounts of purified Sumo-ELL2_519-640_ were incubated for 30 min with 5 nM labeled peptide at room temperature in 25 mM HEPES pH 7.5, 100 mM NaCl, 10% glycerol,0.05% NP40, and 0.5 mM Tris(2-carboxyethyl)phosphine (TCEP) to determine a suitable protein concentration for competition experiments. Competition titration experiments with unlabeled His-tagged AFF4 protein 301-351 were performed using 2 μM Sumo-ELL2 in 25 mM HEPES pH 7.5, 100 mM NaCl, 10% glycerol, 0.05% NP40, 0.5 mM TCEP, and 5 nM fluorescent peptide. Fluorescence anisotropy was measured using a Victor 3V (Perkin Elmer) multi-label plate reader. Data points represent the average of three experiments. Binding curves were fit to a formula describing competitive binding of two different ligands to a protein using Prism version 5.0c (Graphpad Software). Error bars are representative of the standard error from the mean of three experimental replicates.

### Co-immunoprecipitation

Approximately 2 × 10^7^ HEK 293T cells in two 145-mm dishes were transfected by plasmids expressing the wild-type or mutant FLAG-AFF4 or ELL2-HA (20 μg/each). 48 hours after transfection, the cells were harvested and swollen in 4 ml hypotonic buffer A (10 mM HEPES-KOH [pH 7.9], 1.5 mM MgCl_2_, and 10 mM KCl) for 5 minutes and then centrifuged at 362 × g for 5 min. The cells were then disrupted by grinding 20 times with a Dounce tissue homogenizer in 2 ml buffer A, followed by centrifugation at 3,220 × g for 10 min to collect the nuclei. The nuclei were then extracted in 400 μl buffer C (20 mM HEPES-KOH [pH 7.9], 0.42 M NaCl, 25 % glycerol, 0.2 mM EDTA, 1.5 mM MgCl_2_, 0.4 % NP-40, 1 mM dithiothreitol, and 1 × protease inhibitor cocktail) on ice for 30 min, followed by centrifugation at 20,800 × g for 30 min. The supernatant (NE) was then mixed with 10 μl of anti-FLAG agarose (A2220 Sigma) or Anti-HA agarose (A2095 Sigma) and rotated at 4 °C overnight. The beads were then washed three times with buffer D (20 mM HEPES-KOH [pH 7.9], 0.3 M KCl, 15 % glycerol, 0.2 mM EDTA, and 0.4 % NP-40), and eluted with 30 μl 0.1 M glycine-HCl (pH 2.0). For Western blot, 3 % of the NE input and 50 % of the IP eluate were loaded into each NE and IP lane, respectively.

### Luciferase reporter assay

Approximately 6 × 10^5^ HEK 293T cells or 4 × 10^5^ HeLa cells in 6-well plates were transfected in triplicate by plasmids expressing FLAG-AFF4 and/or ELL2-HA (1 μg/each) with the HIV-1 LTR-luciferase construct (0.1 μg). 48 hours after transfection, the cells were harvested and lysed in 1 × Reporter Lysis Buffer (E3971 Promega), followed by centrifugation at 20,800 g for 1 min. Luciferase activities in the supernatant were measured using the Luciferase Assay System (E1501 Promega) on a Lumat LB 9501 luminometer.

## Acknowledgements

We thank Xuefeng Ren, James Holton, and George Meigs for assistance with data collection. This work was supported by NIH grants P50GM082250 (J. H. H.) and NIAID R01AI041757 and R01AI095057 (Q. Z.). The Minstrel crystal farm was purchased with support from the NIH, S10 OD016268. Beamline 8.3.1 at the Advanced Light Source, LBNL, is supported by the U. C. Office of the President, Multicampus Research Programs and Initiatives grant MR-15-328599 and the Program for Breakthrough Biomedical Research, which is partially funded by the Sandler Foundation. The Advanced Light Source is supported by the Director, Office of Science, Office of Basic Energy Sciences, of the U.S. Department of Energy under Contract No. DE-AC02-05CH11231. The Stanford Synchrotron Radiation Lightsource is supported by the U. S. D. O. E. under contract No. DE-AC02-76SF00515. The SSRL Structural Molecular Biology Program is supported by the DOE Office of Biological and Environmental Research and by NIH grant P41GM103393. Coordinates have been deposited in the RCSB.

## Figure Legends

**Figure 1 Supplement 1.**
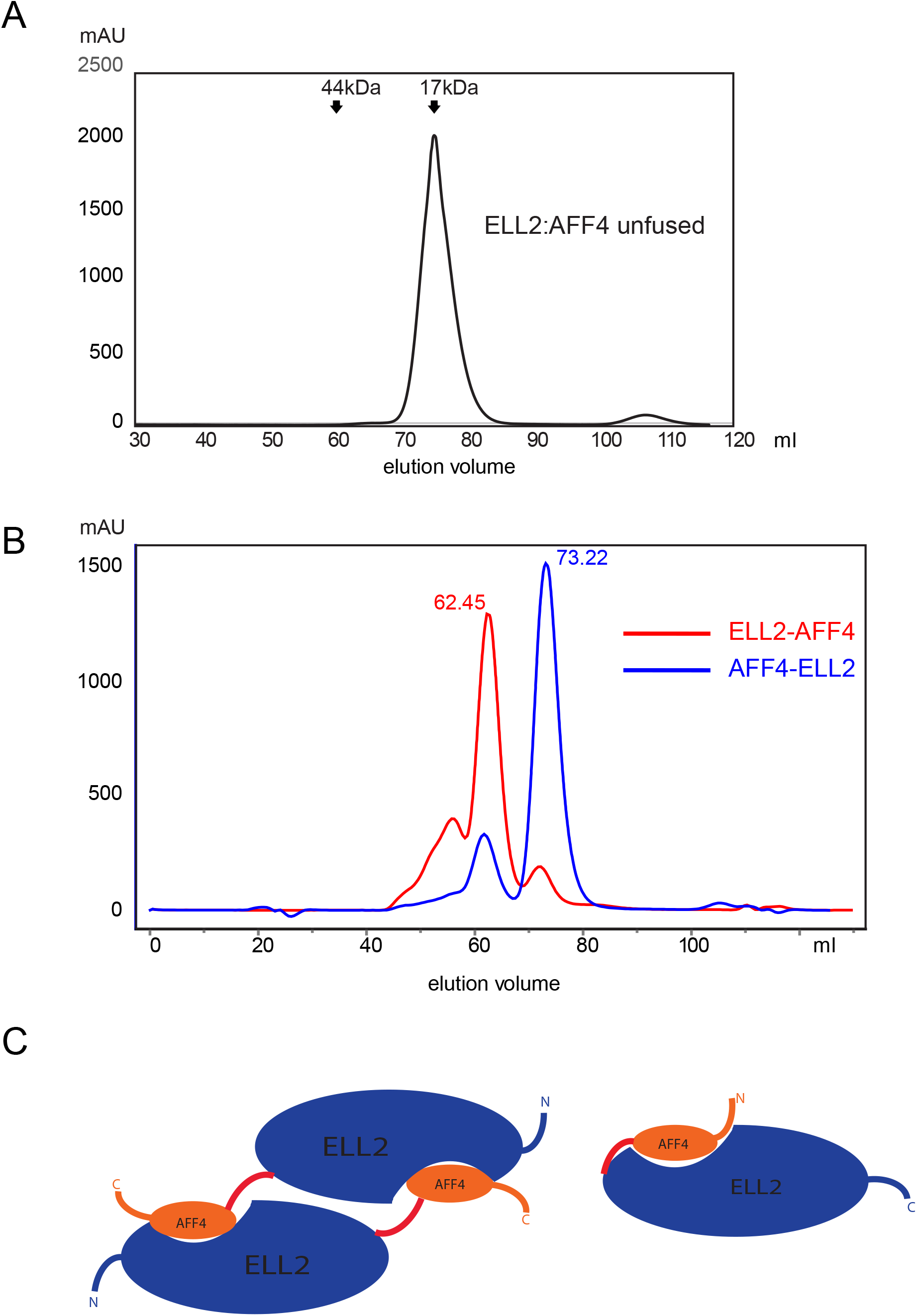
Fusion constructs screen. **A.** The unfused ELL2^Occ^:AFF4^ELLBow^ complex is monomeric in solution. **B.** The ELL2^Occ^–(Gly-Ser)4-AFF4^ELLBow^ was eluted at 62.45ml on the Hiload 16/60(GE) while AFF4^ELLBow^–(Gly-Ser)_4_-ELL2^Occ^ was eluted at 73.22 ml, which correspond to a dimer and a monomer, respectively. Red line: ELL2^Occ^–(Gly-Ser)4-AFF4^ELLBow^, Blue line: AFF4^ELLBow^–(Gly-Ser)4-ELL2^Occ^. **C.** Schematic of hypothesis for ELL2^Occ^–(Gly-Ser)_4_-AFF4^ELLBow^ dimerization in solution while AFF4^ELLBow^–(Gly-Ser)_4_-ELL2^Occ^ was monomeric. ELL2^Occ^ is shown in blue. AFF4^ELLBow^ is shown in orange. N, C represent amino termini and carboxyl termini respectively.

**Figure 2 Supplement 1.**
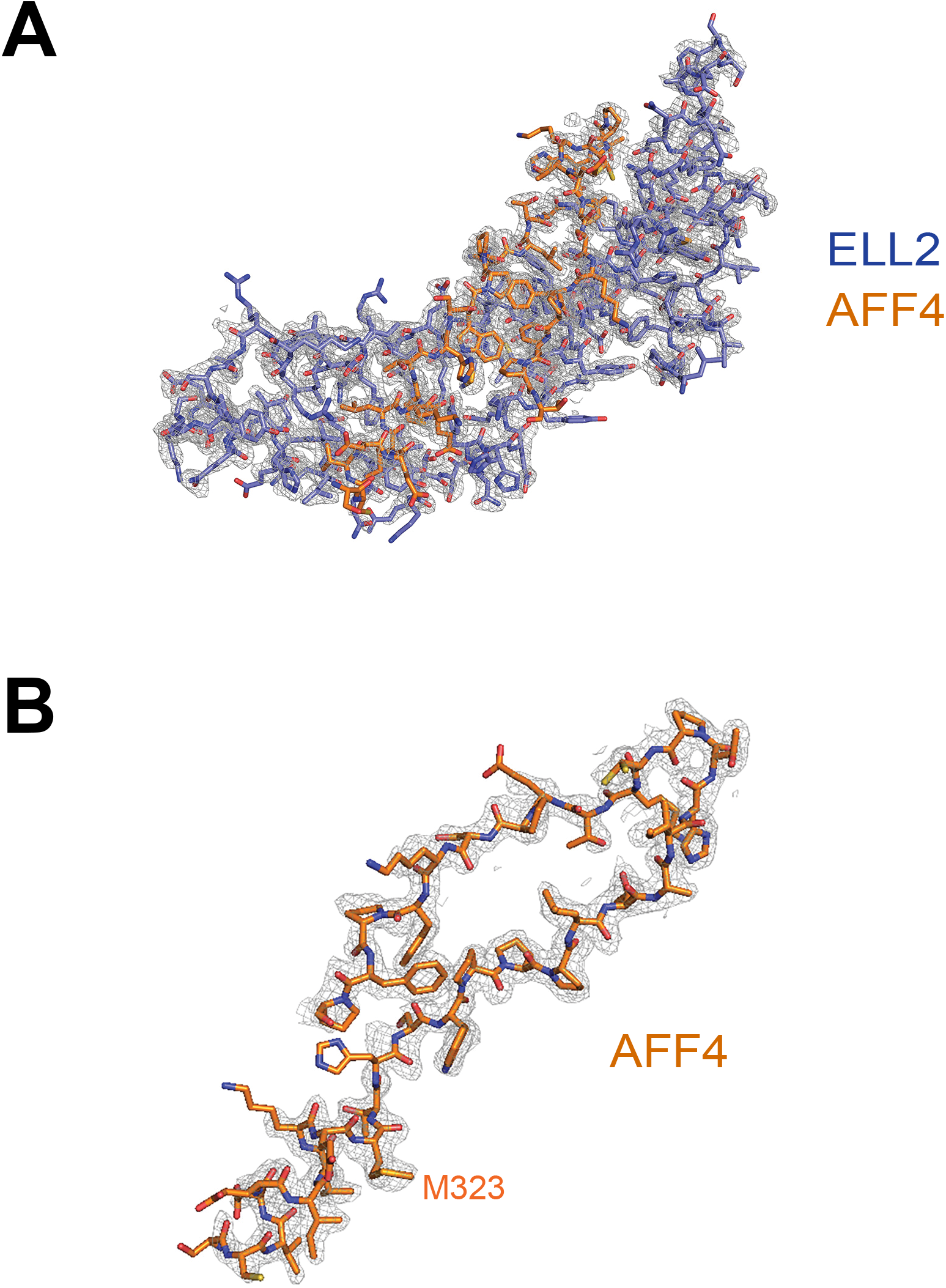
Electron density. **A.** The overall experimental electron density map after density modification is displayed at a contour level of 2σ (gray), with AFF4^ELLBow^ and ELL2^Occ^ shown as stick in orange and light blue, respectively. **B.** The map from (A) corresponding to AFF4^ELLBow^ is displayed at a contour level of 2σ (gray), with AFF4^ELLBow^ shown in a stick model. Met323 is highlighted.

**Figure 7 Supplement 1.**
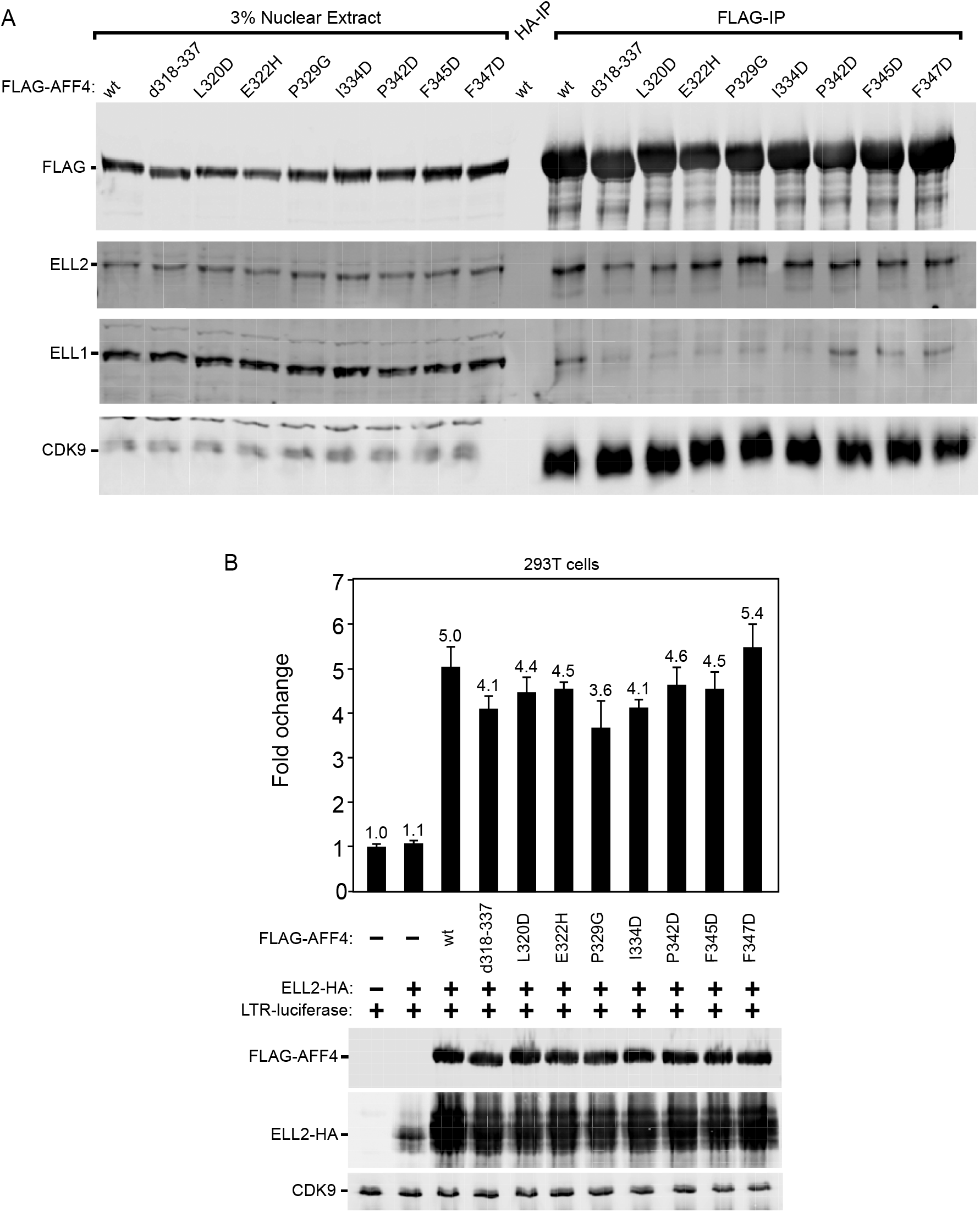
AFF4 ELLBow mutants in the background of the full-length protein have little effect on binding or proviral transcription. **A.** NE and IP were conducted as in Fig. 7 using HEK 293T cells transfected by the indicated plasmids. **B.** Luciferase activities were measured and analyzed as in Fig. 7 using extracts of HEK 293T cells transfected with the combinations of the indicated plasmids.

